# Transcriptomic response to histone deacetylase inhibitors in cultured feline cells

**DOI:** 10.64898/2026.06.08.731028

**Authors:** Ganma Tanaka, Shizune Nakamura, Rikuto Goto, Akiko Kubota, Naoaki Sakamoto, Akinori Awazu

**Affiliations:** Graduate School of Integrated Sciences for Life, Hiroshima University, Higashi-hiroshima, Hiroshima, Japan; Research Center for the Mathematics on Chromatin Live Dynamics, Hiroshima University, Higashi-hiroshima, Hiroshima, Japan

## Abstract

**Objective:** In recent years, the number of cats kept as companion animals has increased, leading to a growing demand for veterinary care. Although some histone deacetylase (HDAC) inhibitors are promising for the treatment of human cancers and neurological diseases, comprehensive systematic research on HDAC inhibitors in domestic cats remains insufficient. Therefore, this study aimed to investigate the effects of HDAC inhibitors on the transcriptome of feline cells.

**Methods:** Two types of cells derived from domestic cats, Crandell-Rees Feline Kidney (CRFK; kidney-derived) cells and PG-4 cells (astrocyte-derived), were treated with four HDAC inhibitors (panobinostat, trichostatin A, valproic acid, and vorinostat) for 24 h. Transcriptomic changes after treatment were examined using RNA sequencing.

**Results:** HDAC inhibitor treatment upregulated the expression of intercellular chemical interactions and signal transduction-related genes, similar to observations in human cells. Although HDAC inhibitors did not suppress the expression of cell cycle-related genes in CRFK cells, as observed in human cells, the inhibitors downregulated the expression of organogenesis-related genes. Consistent with observations in human cells, HDAC inhibitors suppressed the expression of cell cycle- and cancer-related genes in PG-4 cells. Importantly, valproic acid, which is thought to be more effective for neurological diseases than for cancer, suppressed the expression of more cancer-related genes in PG-4 cells than the other three HDAC inhibitors.

**Conclusion and relevance:** Our findings revealed that the responses of cells derived from feline organs to various HDAC inhibitors varied considerably depending on the organ of origin and species. Since few studies, including human studies, have comprehensively compared transcriptomic responses to multiple HDAC inhibitor classes across multiple cell types, the results of this study provide a foundation for future research on the treatment and prevention of cancer and neurological diseases in domestic cats and other mammals.

## Introduction

Histone deacetylases (HDACs) are enzymes that play a central role in epigenetic gene regulation by promoting histone deacetylation, altering chromatin structure, and repressing the transcription of specific genes.^1^ Previous studies have shown that HDAC inhibitors affect various gene expression pathways.^2–4^ Notably, these inhibitors have been widely studied in human cancer, cardiovascular disease, neurodegenerative diseases, inflammatory diseases, autoimmune diseases, metabolic diseases, and anti-aging.^5–11^

Domestic cats (*Felis catus*) exhibit diseases similar to those in humans. Over the years, the efficacy of HDAC inhibitors has been investigated for the treatment of mammary gland cancer, feline immunodeficiency virus (FIV) infection, and cardiopulmonary disease.^12–14^ Treatment with HDAC inhibitors has been shown to induce apoptosis in a feline mammary gland cancer cell line and improve function in a feline cardiopulmonary model.^12,13^ In addition, the HDAC inhibitor vorinostat (VO; suberoylanilide hydroxamic acid) has demonstrated therapeutic efficacy in an FIV-infected cat model^14^ and has been clinically effective in the treatment of other diseases. ^15^

Notably, the FDA has approved VO, panobinostat (PA), and valproic acid (VA) for the treatment of various human diseases.^5,16–19^ For instance, VO was approved by the FDA in 2006 for the treatment of cutaneous manifestations in patients with cutaneous T-cell lymphoma (CTCL).^20^ PA was approved by the FDA in 2015 for use in combination with bortezomib and dexamethasone in patients with multiple myeloma (but has now been withdrawn). In addition, VA was approved by the FDA for the treatment of epilepsy (1978), bipolar disorder (1995), and migraine prophylaxis (2008).^21–23^ Although TS has not been approved by the FDA, it is known for its anticancer properties and biphasic inflammatory response.^24–25^

However, a comprehensive analysis of gene expression changes following the administration of various HDAC inhibitors in domestic cats has not been performed. In addition, the differences in response between inhibitors and cat tissue-specific response patterns remain unclear. To address this gap, this study aimed to investigate the effects of four HDAC inhibitors (PA, TS, VA, and VO) on the transcriptome of two cultured cell lines derived from domestic cats, Crandell-Rees Feline Kidney (CRFK; kidney-derived) cells and PG-4 cells (astrocyte-derived), and comprehensively examine changes in gene expression using RNA-seq. To the best of our knowledge, this study is the first to comprehensively analyze the transcriptomic response of feline cells to HDAC inhibitors. Overall, this study is anticipated to provide fundamental data for examining the clinical efficacy of HDAC inhibitor drugs in the treatment of feline diseases.

## Materials and methods

### Cell lines and culture conditions

CRFK cells (JCRB9035, RRID: CVCL_2426) and feline sarcoma-positive leukemia-negative (S+L−) astrocyte cells termed PG-4 (S+L-) (JCRB9125, RRID: CVCL_3322) were obtained from the JCRB Cell Bank (National Institutes of Biomedical Innovation, Health and Nutrition, Osaka, Japan). CRFK cells were cultured in Dulbecco’s Modified Eagle Medium (DMEM; Gibco, Thermo Fisher Scientific, Waltham, MA, USA) supplemented with 10% fetal bovine serum (FBS), penicillin (100 IU/mL), streptomycin (100 ng/mL) (Sigma-Aldrich, St. Louis, MO, USA), and 1% non-essential amino acids (FUJIFILM Wako Pure Chemical Corporation, Osaka, Japan) at 37 °C under 5% CO_2_ conditions. PG-4 cells were cultured in McCoy’s 5A Modified Medium (Gibco) supplemented with 10% heat-inactivated FBS, penicillin (100 IU/mL), and streptomycin (100 ng/mL) at 37 °C under 5% CO_2_ conditions. For cell passaging, the cells were washed using BASIC DPBS (no calcium or magnesium; Gibco) and treated with TrypLE™ Express Enzyme (1X; Gibco) at 37 °C for 5 min under 5% CO_2_ conditions.

### Treatment of cultured cells with HDAC inhibitors

Briefly, cultured cells were treated with the following four HDAC inhibitors: PA (SML3060-10MG: Sigma-Aldrich), TS (T1952-200UL: Sigma-Aldrich), VA (1708707-500MG: Sigma-Aldrich), and VS (SML0061-5ML: Sigma-Aldrich). Specifically, cells were seeded in 35 mm tissue culture dishes with a polystyrene coating containing 2 ml of the same medium used for culture until the cell density reached 60–80% confluence. After 24 h, the culture medium was removed and replaced with 2 ml of medium containing specific concentrations of an HDAC inhibitor. Notably, treatment with the inhibitors was performed as follows: 1 ml of culture medium without an HDAC inhibitor was first added to the dish, and then 1 ml of culture medium containing a double concentration of the HDAC inhibitor was added to make 2 ml of culture medium with the desired HDAC inhibitor concentration.

In addition, the HDAC inhibitor concentrations in the medium were selected such that the net viable cell count (cells attached to the bottom of the dish) at 0 and 24 h of culture did not exhibit large variations, as confirmed in three replicated experiments (Table S1). In particular, the concentrations were as follows: PA, 0.056 nM for CRFK and 0.260 nM for PG-4; TS, 0.156 nM for CRFK and 0.556 nM for PG-4; VA, 9.360 µM for CRFK and 15.563 µM for PG-4; and VO, 2.58 nM for CRFK and 23.23 nM for PG-4. Cell counts were performed using a LUNA™ Automated Cell Counter (Logos Biosystems, Gyeonggi-do, South Korea) and LUNA™ Cell Counting slides (Logos Biosystems). Cells that detached from the bottom of the dish and floated were considered dead and counted as dead cells.

In this study, cell samples before HDAC inhibitor administration (0 h) and 24 h after treatment served as the control and treatment groups, respectively. As described above, the cell density was kept constant between the control and HDAC inhibitor-treated samples. Therefore, we could evaluate the differences in HDAC inhibitor-dependent gene expression changes under nearly identical conditions for cell density and net inhibitor toxicity across the samples.

### RNA sequencing

Total RNA was extracted from the control and treated samples using ISOGEN (NIPPON GENE, Tokyo, Japan) according to the manufacturer’s protocol. Thereafter, the extracted RNA was treated with recombinant DNase I (RNase-free) (TaKaRa Bio, Japan) and purified using an RNeasy Mini Kit (QIAGEN, Hilden, Germany) according to the manufacturer’s protocol.

RNA sequencing was performed by Novogene Corporation (Beijing, China). For library preparation, mRNA was enriched using poly-T oligo-attached magnetic beads to select poly(A)+ transcripts. Strand-specific cDNA libraries were constructed using the NEBNext® Ultra™ RNA Library Prep Kit for Illumina® (New England Biolabs, Ipswich, MA, USA), incorporating dUTP during second-strand synthesis to preserve strand information. Libraries were sequenced on an Illumina NovaSeq X Plus platform using a paired-end 150 bp strategy, generating approximately 6 Gb of raw data per sample, which were obtained as .fastq files.

Two biological replicates of RNA-seq data were obtained for the control and each treatment sample.The RNA integrity number (RIN) of all extracted RNA for RNA-seq was > 9.6 (Table S2).

### Quantification of RNA sequencing data

Quality control of each raw read in each .fastq file was performed using fastp (version 0.22.0) with the default parameters.^26^ Filtered reads were aligned to the reference genome sequence of *Felis catus*, F.catus_Fca126_mat1.0 (RefSeq assembly GCF_018350175.1), using STAR (version 2.7.4a) with the options -outSAMtype BAM SortedByCoordinate, -quantMode TranscriptomeSAM, -outSAMattributes All, -outFilterMultimapNmax 1000, -outSAMunmapped Within KeepPairs, and -outSAMstrandField intronMotif.^27^ Transcript quantification as count data was performed using RSEM (version 1.3.1) with the options -estimate-rspd, -strandedness reverse, and -no-bam-output.^28^

### Analysis of quantified transcriptome data

To ensure the reproducibility of the results, the quantified transcriptome data were analyzed using iDEP v2.4.4 (https://bioinformatics.sdstate.edu/idep/).^29^ Differential expression analysis was performed to identify differentially expressed genes (DEGs) between each HDAC inhibitor group and the control group using the Wald test with default parameters (FDR cutoff = 0.1 and minimum fold-change = 2) using read count data of RNA-seq data (results are shown in Table S3.). Gene ontology (GO) functional annotation and Kyoto Encyclopedia of Genes and Genomes (KEGG) pathway enrichment analyses of the DEGs was performed using the “Stats” and “enrichment” functions in “DEG” function of iDEP v2.4.4. GO terms with FDR < 10^-5^ and KEGG pathways with FDR < 0.1 were considered significant terms and pathways, respectively.

## Results

### DEGs following treatment with each HDAC inhibitor

In this study, we identified DEGs between each HDAC group (PA, TS, VA, and VO) and the control group. Notably, more than 2000 significantly upregulated and 1000 significantly downregulated genes were identified in CRFK and PG-4 cells following HDAC inhibitor treatment (Figure 1; Table S3). In addition, the number of significantly upregulated and downregulated genes was higher in PG-4 cells than in CRFK cells.

**Figure 1.**
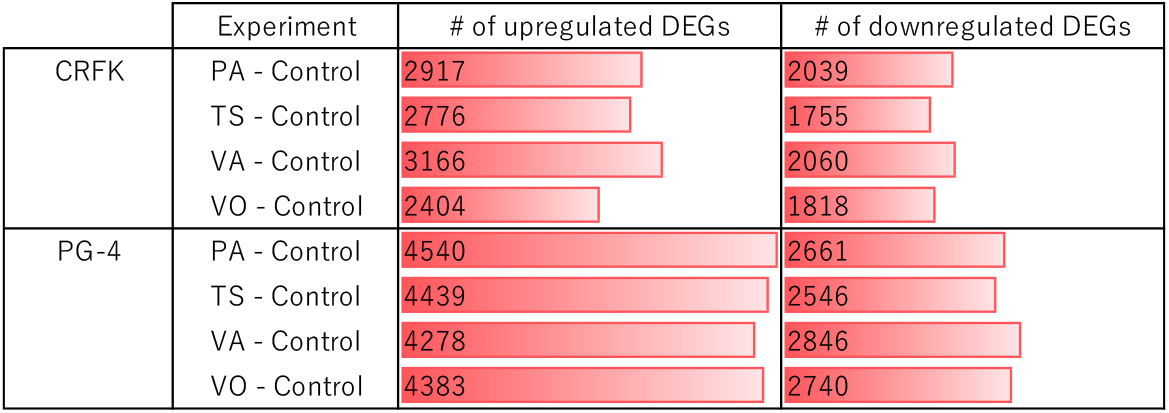
Number of differentially expressed genes (DEGs) in CRFK and PG-4 cells treated with various HDAC inhibitors. Number of upregulated and downregulated DEGs in CRFK and PG-4 cells treated with panobinostat (PA), trichostatin A (TS), valproic acid (VA), and vorinostat (VO). Each value and bar length indicate the number of DEGs in the treated sample compared with the control.

### Common features of DEGs in response to HDAC inhibitors

GO and KEGG pathway enrichment analyses of DEGs were performed to identify the functional terms and pathways affected by HDAC inhibitors. Treatment with the four HDAC inhibitors upregulated genes enriched in “System process,” “Multicellular organismal process,” and “Nervous system process” in the GO database (Figure 2) and “Neuroactive ligand-receptor interaction” in the KEGG database in CRFK and PG-4 cells (Figure 3). In addition, treatment with the inhibitors upregulated genes related to intercellular communication, including those involved in “Cell-cell signaling,” “Synaptic signaling,” and “Monoatomic ion transmembrane transport” in the GO database and “Nicotine addiction” and “Calcium signaling pathway” in the KEGG database (except the response to VO in the CRFK cell) in both cells (Figure 2 and 3). Moreover, treatment with the inhibitors upregulated genes involved in the “Synaptic vesicle cycle,” “Circadian entrainment,” and “Serotonergic synapse” in the KEGG database (except for the response to VO in the CRFK cell and VA in the PG-4 cell) in both cell lines (Figure 2 and 3).

**Figure 2.**
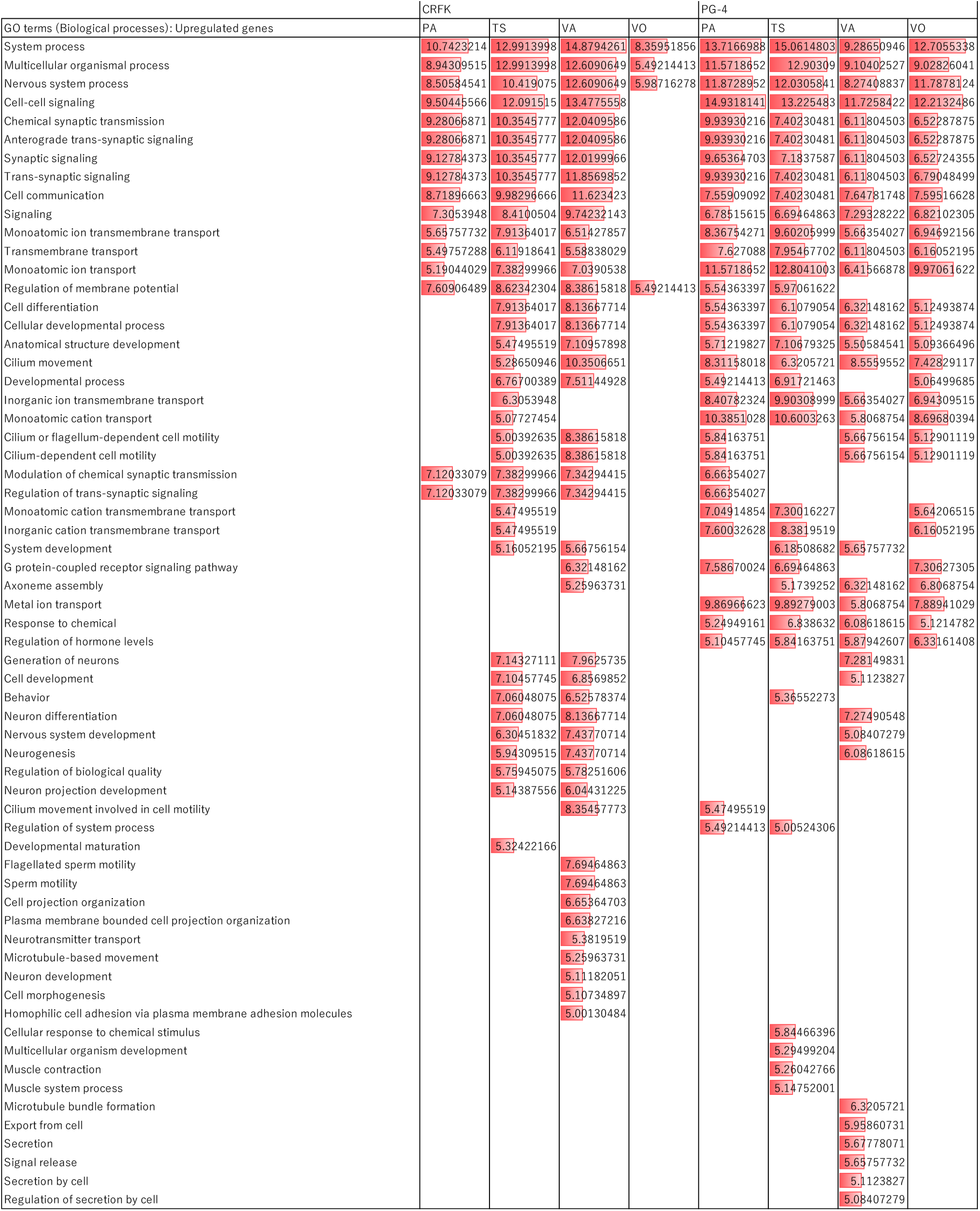
Gene ontology (GO) biological processes significantly enriched by upregulated genes by each HDAC inhibitor in CRFK and PG-4 cells. Each score and red bar length indicate the−log_10_(FDR) value.

**Figure 3.**
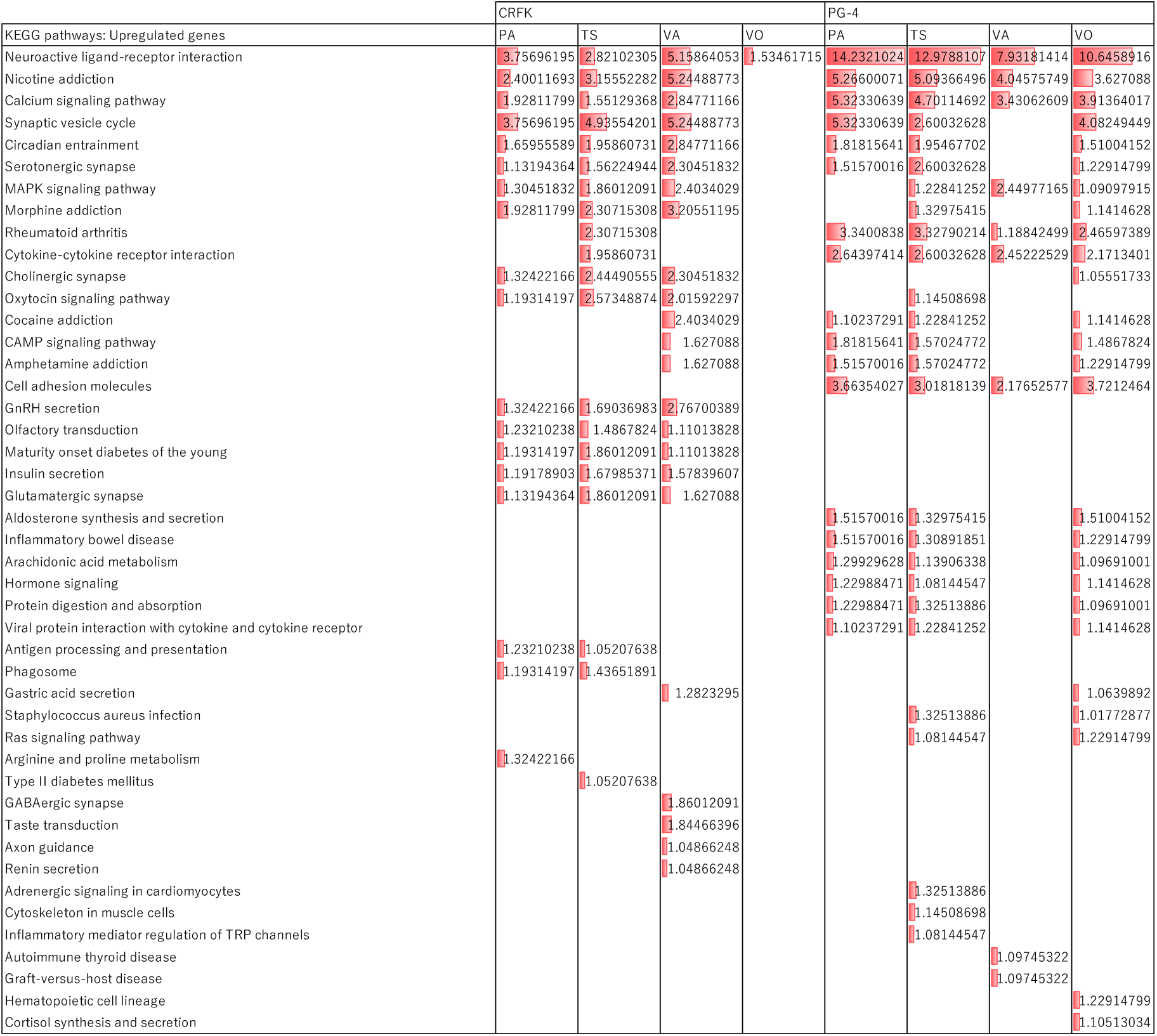
Kyoto Encyclopedia of Genes and Genomes (KEGG) pathways significantly enriched by upregulated genes by each HDAC inhibitor in CRFK and PG-4 cells. Each score and red bar length indicate −log_10_(FDR) value

In contrast, no common features of downregulated genes in response to the four HDAC inhibitors in the two cell lines were obtained, and only “Negative regulation of cellular biosynthetic process” in the GO database was affected by the four HDAC inhibitors, except for the response to TS in the CRFK cell (Figure 4–6).

**Figure 4.**
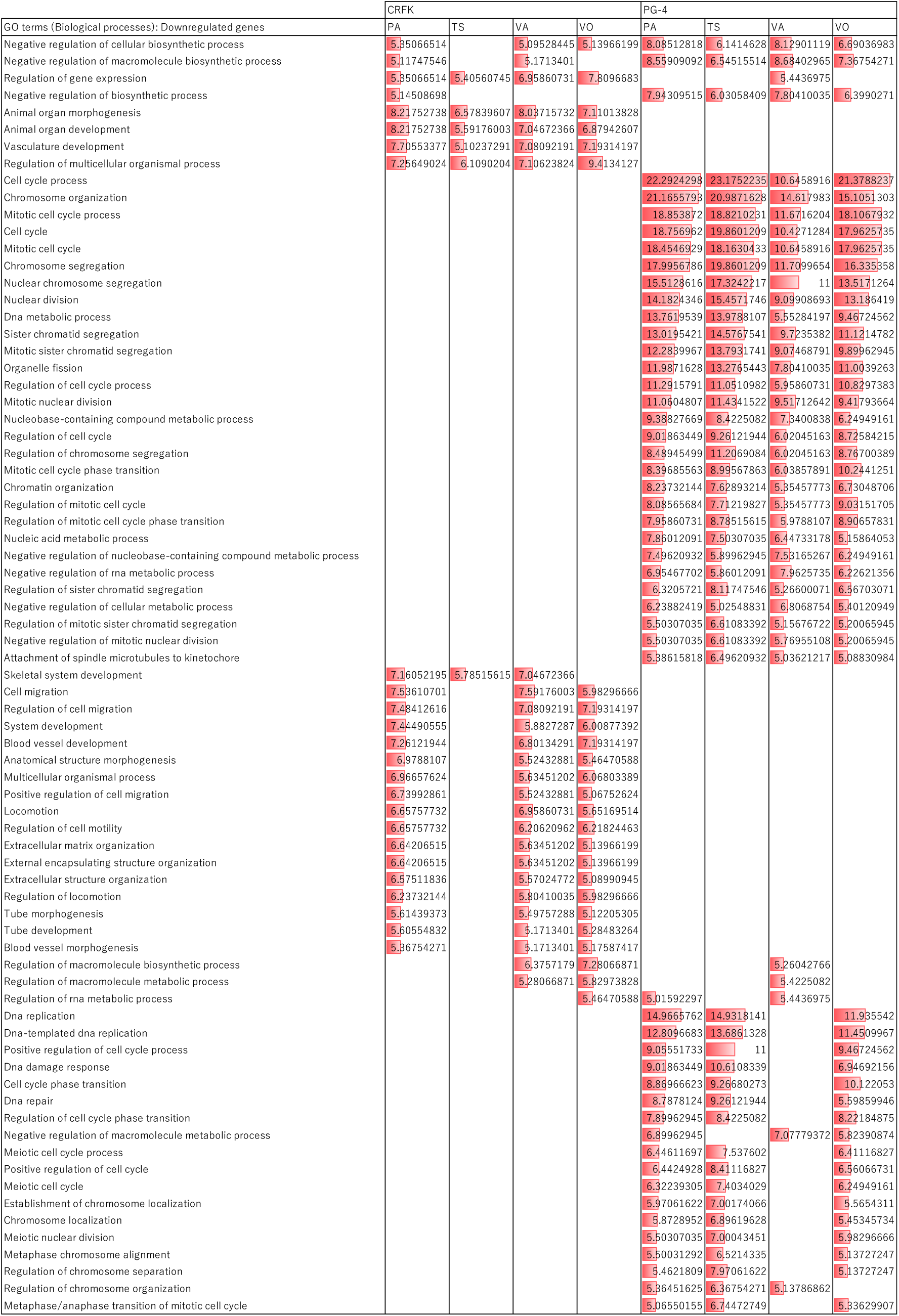
Gene ontology (GO) biological processes significantly enriched by downregulated genes by each HDAC inhibitor in CRFK and PG-4 cells, with GO terms identified in at least three combinations of cell types and HDAC inhibitors listed. Each score and red bar length indicate the −log_10_(FDR) value.

**Figure 5.**
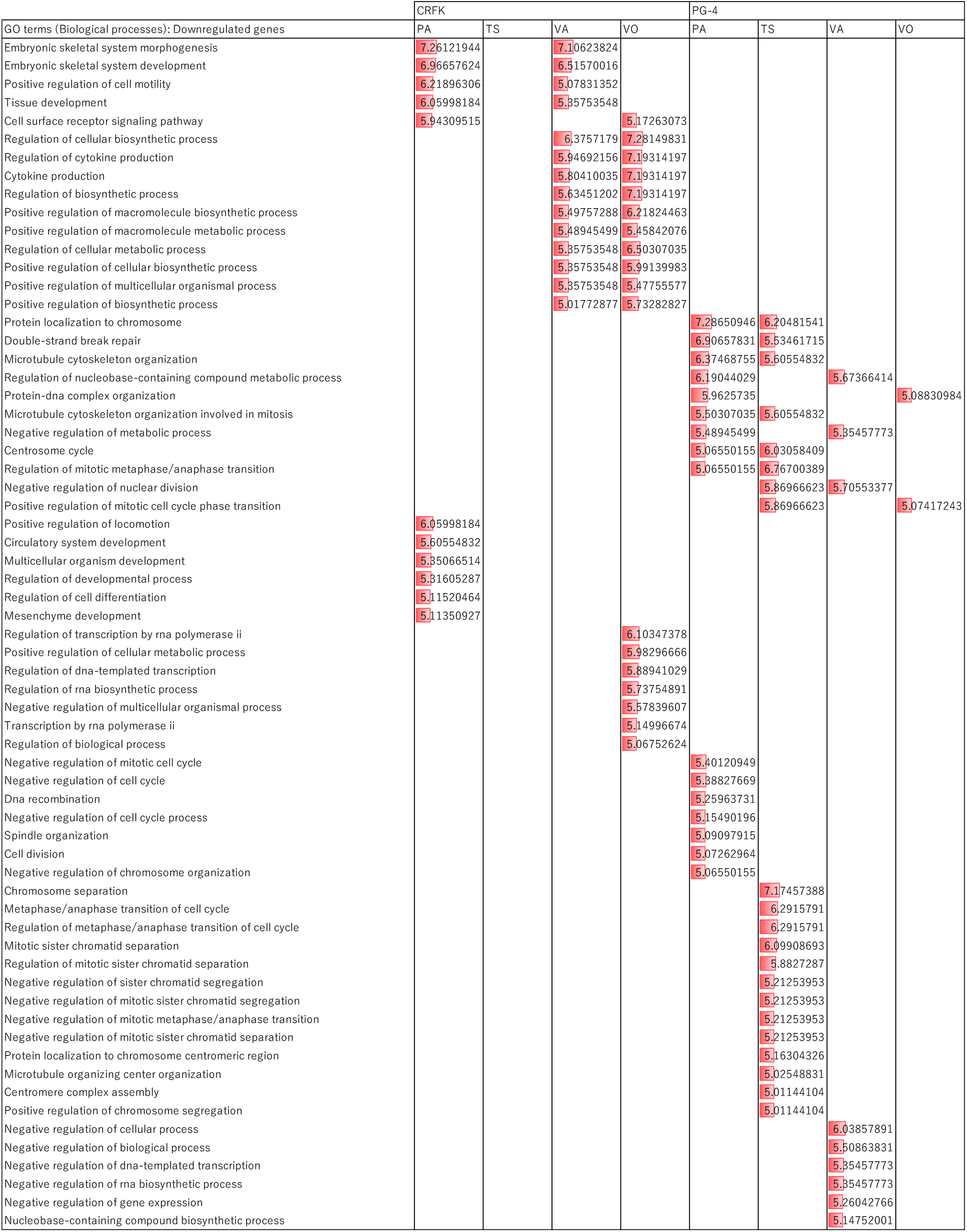
Gene ontology (GO) biological processes significantly enriched by downregulated genes by each HDAC inhibitor in CRFK and PG-4 cells, with GO terms identified in at least one combination of cell types and HDAC inhibitors listed. Each score and red bar length indicate the −log_10_(FDR) value.

**Figure 6.**
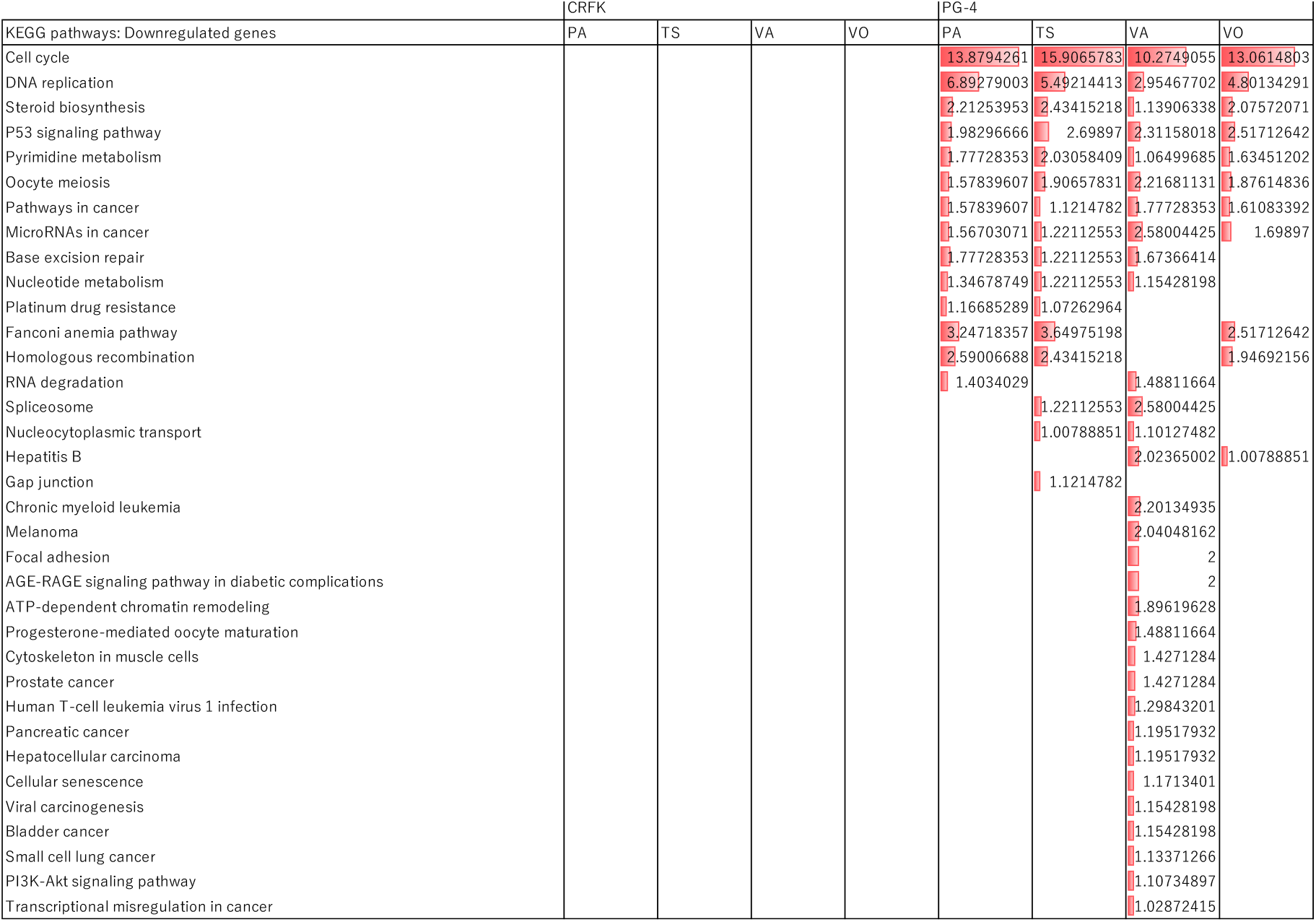
Kyoto Encyclopedia of Genes and Genomes (KEGG) pathways containing a significantly large number of downregulated genes by each HDAC inhibitor in CRFK and PG-4 cells. Each score and red bar length indicate the −log_10_(FDR) value.

### Cell-specific features of upregulated genes in response to HDAC inhibitors

Treatment with HDAC inhibitors upregulated genes related to the GO term “Regulation of membrane potential” in CRFK cells (Figure 2 and 3). In addition, treatment with the inhibitors upregulated genes related to intercellular chemical communications and signaling processes, including those involved in “Modulation of chemical synaptic transmission” and “Regulation of trans-synaptic signaling” in the GO database, and “MAPK signaling pathway,” “Morphine addiction,” “Cholinergic synapse,” and “Oxytocin signaling pathway” in the KEGG database (except the response to VO) in CRFK cells (Figure 2 and 3).

In contrast, PG-4 cells exhibited the upregulation of genes related to cell differentiation, multicellular structure development, and intercellular communications, including those involved in “Cell differentiation,” “Anatomical structure development,” “Cilium movement,” and “Regulation of hormone levels” in the GO database and “Cytokine-cytokine receptor interaction” and “Cell adhesion molecules” in the KEGG database in response to the four HDAC inhibitors (Figure 2 and 3). In addition, treatment with HDAC inhibitors upregulated genes related to intercellular chemical communication, including “Developmental process,” “Monoatomic cation transmembrane transport,” and “G protein-coupled receptor signaling pathway,” in the GO database and “CAMP signaling pathway,” “Hormone signaling,” and “Protein digestion and absorption” in the KEGG database (except the response to VA) in PG-4 cells (Figure 2 and 3).

### Cell-specific features of downregulated genes set in response to HDAC inhibitors

Treatment with the four HDAC inhibitors downregulated genes related to the GO terms “Animal organ morphogenesis,” “Animal organ development,” “Vasculature development,” and “Regulation of multicellular organismal process” in CRFK cells (Figure 4). In addition, CRFK cells exhibited downregulation of genes related to cell migration and blood vessel development, including those involved in “Cell migration,” “Blood vessel development,” “Extracellular structure organization,” and “Tube morphogenesis” in the GO database (except for the response to TS) following treatment with HDAC inhibitors (Figure 4).

In contrast, PG-4 cells exhibited the downregulation of genes related to cell cycle, and chromosome segregation, and cancer-related processes, including those involved in “Cell cycle process,” “Chromosome segregation,” and “Mitotic nuclear division” in the GO database, and “Cell cycle,” “DNA replication,” “P53 signaling pathway,” and “Pathways in cancer” in the KEGG database following treatment with the four HDAC inhibitors (Figure 4 and 6).

### HDAC inhibitor-specific response features in CRFK cells

CRFK cells exhibited upregulation of genes related to intercellular communication, including those involved in “Cilium movement,” “Generation of neurons,” and “Neurotransmitter transport” in the GO database and “GABAergic synapse,” “Taste transduction,” and “Axon guidance” in the KEGG database following VA treatment (Figure 2 and 3). CRFK cells also exhibited a partly similar response to TS treatment to that of VA treatment.

In addition, CRFK cells exhibited downregulation of genes related to the organization of multicellular structures, including those involved in “Embryonic skeletal system morphogenesis,” “Tissue development,” “Circulatory system development,” and “Mesenchyme development” in the GO database following PA treatment (Figure 4 and 6). Moreover, CRFK cells exhibited downregulation of genes related to intracellular metabolic and transcriptional processes, including those involved in “Positive regulation of macromolecule biosynthetic process,” “Regulation of cellular metabolic process,” and “Regulation of transcription by RNA polymerase II” in the GO database following VO treatment (Figure 4 and 5). CRFK cells also showed a partly similar response to VA treatment to that of PA and VO treatments. In contrast, CRFK cells exhibited minimal downregulation of genes related to specific physiological processes in both GO and KEGG databases following TS treatment (Figure 5 and 6).

### HDAC inhibitor-specific response features in PG-4 cells

PG-4 cells exhibited further upregulation of genes related to signal molecule secretion, including those involved in “Export from cell,” “Signal release,” and “Regulation of secretion by cell” in the GO database after VA treatment, even though PG-4 cells exhibited slight upregulation of genes related to specific physiological processes in the KEGG database (Figure 2 and 3).

PG-4 cells exhibited further downregulation of genes related to the negative regulation of transcription and pathways in various cancers, such as those involved in “Negative regulation of DNA-templated transcription” and “Negative regulation of RNA biosynthetic process” in the GO database following VA treatment. In addition, KEGG analysis revealed that VA treatment upregulated genes related to “Chronic myeloid leukemia,” “Melanoma,” “Prostate cancer,” and “Pancreatic cancer.” Moreover, PG-4 cells showed modest downregulation of genes associated with the cell cycle in the VA group compared to other HDAC inhibitor groups (Figure 4–6).

## Discussions

Recently, HDAC inhibitors have been shown to induce the following responses in several human cells, although the details may differ depending on the cell type. Generally, HDAC inhibitors increase the expression of genes related to intercellular interactions, signaling pathways, and ion and small molecule transport, which are typically expressed in the nervous system, as well as cell differentiation and apoptosis-related genes, and decrease those that drive the cell cycle, as well as genes related to tumorigenesis, cancer progression, and extracellular structure formation.^30–37^ Previous studies have investigated the potential anticancer effects of PA, TS, and VO in humans.^5,16–20,24–25^ VA has a slightly different specificity than the other three HDAC inhibitors and has mainly attracted attention as a treatment for neurological disorders. ^21–23^

In this study, CRFK and PG-4 cells (derived from domestic cats) exhibited increased expression of genes involved in intercellular interactions, similar to human cells, particularly nerve cells. Notably, CRFK cells tended to show increased expression of genes involved in a wider range of signaling pathways than PG-4 cells. In contrast, PG-4 cells exhibited increased expression of genes involved in cell differentiation, morphogenesis, and ion and small molecule transport.

Contrary to the findings for PG-4 genes, the expression of cell cycle-related genes was not suppressed in CRFK cells. Except for TS, the other HDAC inhibitors decreased the expression of organogenesis-related genes in CRFK cells. Consistent with observations in human cells, PG-4 cells exhibited a decrease in the expression of genes involved in cell cycle, cell proliferation, cancer-related pathways, and p53-related pathways. Notably, VA treatment affected different pathways in PG-4 cells compared with the other three HDAC inhibitors. Treatment with the other three HDAC inhibitors reduced the expression of genes related to the cell cycle, DNA replication, chromosome segregation, and cell division. In contrast, VA treatment reduced the expression of genes involved in pathways that negatively regulate transcription and tumorigenesis. Importantly, while VA is thought to be more effective for neurological diseases than for cancer in human medicine, VA suppressed the expression of more cancer-related genes in PG-4 cells than the other three HDAC inhibitors; the FDA has approved VO and PA for the treatment of tumor-related manifestations.

Notably, neither cell type exhibited increased expression of apoptosis-related genes following treatment with HDAC inhibitors, contrary to findings in human cells and feline mammary cancer cell lines.^12^ However, this may be due to the concentration of HDAC inhibitors being lower than that used in typical clinical studies. Therefore, increasing the concentration of HDAC inhibitors may increase the expression of apoptosis-related genes.

In this study, we compared the responses of cells from different organs of domestic cats to various HDAC inhibitors. To the best of our knowledge, this is the first comprehensive analysis of the transcriptomic response to HDAC inhibitors in feline cells. Although there have been a few reports of microarray studies on human cells,^2,30^ no studies, including those on human cells, have comprehensively compared the transcriptomic responses of multiple cell types to various HDAC inhibitors using RNA-seq. RNA-seq allows for highly flexible analyses, such as the quantitative determination of isoform expression. Importantly, our study provides a theoretical foundation for future research on the treatment and prevention of cancer and neurological diseases in domestic cats and other mammals, including humans.

Despite these promising findings, our study had some limitations that should be considered. First, in this study, the types and concentrations of cells used in the experiment, as well as the types, concentrations, and administration methods of HDAC inhibitors, were limited. For instance, the cells used for comparison in this study were only astrocyte-derived cells (PG-4) from the fetus of an individual with a white spotted coat and kidney-derived cells (CRFK) from an individual without a white coat.^38^ The presence or absence of white spots in the coat is due to differences in the expression dynamics of the KIT gene.^39^ The KIT gene is a proto-oncogene, and its isoform expression ratio also plays an important role in the development of pigment and the nervous system.^39–40^ Therefore, it is unclear whether the differences in the results between CRFK and PG-4 in this study are due to differences in the organs from which the cells originated or to differences in the expression dynamics of KIT isoforms associated with cancer and neurological diseases. In addition, experiments were conducted only using HDAC inhibitor concentrations that neither significantly increased nor decreased the net cell count to observe the cellular response to HDAC inhibitors. However, this concentration is approximately 30–1000 times lower than that used in conventional human clinical studies examining the effects on tumor cells and side effects on normal cells. To facilitate the clinical application of these findings, the response to higher concentrations of HDAC inhibitors should be investigated. Therefore, future studies should conduct experiments with various cell types and HDAC inhibitors under different conditions and compare the results.

Second, this paper did not focus on the detailed mechanisms of gene expression changes and alterations in histone modification distribution induced by each HDAC inhibitor. This remains a subject for future research.

Third, since this study was *in vitro, in vivo* experiments are also necessary to validate our findings.

---

**Table S1** Concentrations of histone deacetylase (HDAC) inhibitors in the culture medium administered to each cell type, the number of viable cells before HDAC inhibitor administration (0 h), and the number of viable and dead cells 24 h after HDAC inhibitor treatment, in a 35 mm tissue culture dish containing 2 ml of the medium, obtained from three replicate experiments using CRFK cells and PG-4 cells.

**Table S2** List of total RNA volume and RNA integrity numbers (RINs) of RNA solutions used for RNA sequencing, extracted from two replicates of CRFK and PG-4 cells: a control sample and an HDAC inhibitor-treated sample.

**Table S3** Gene list and Wald test results for DESeq2 implemented in iDEP v2.4.4.

